## Supplemental Figures for "Cytotoxic CD4^+^ T cells driven by T-cell intrinsic IL-18R/MyD88 signaling predominantly infiltrate *Trypanosoma cruzi*-infected hearts"

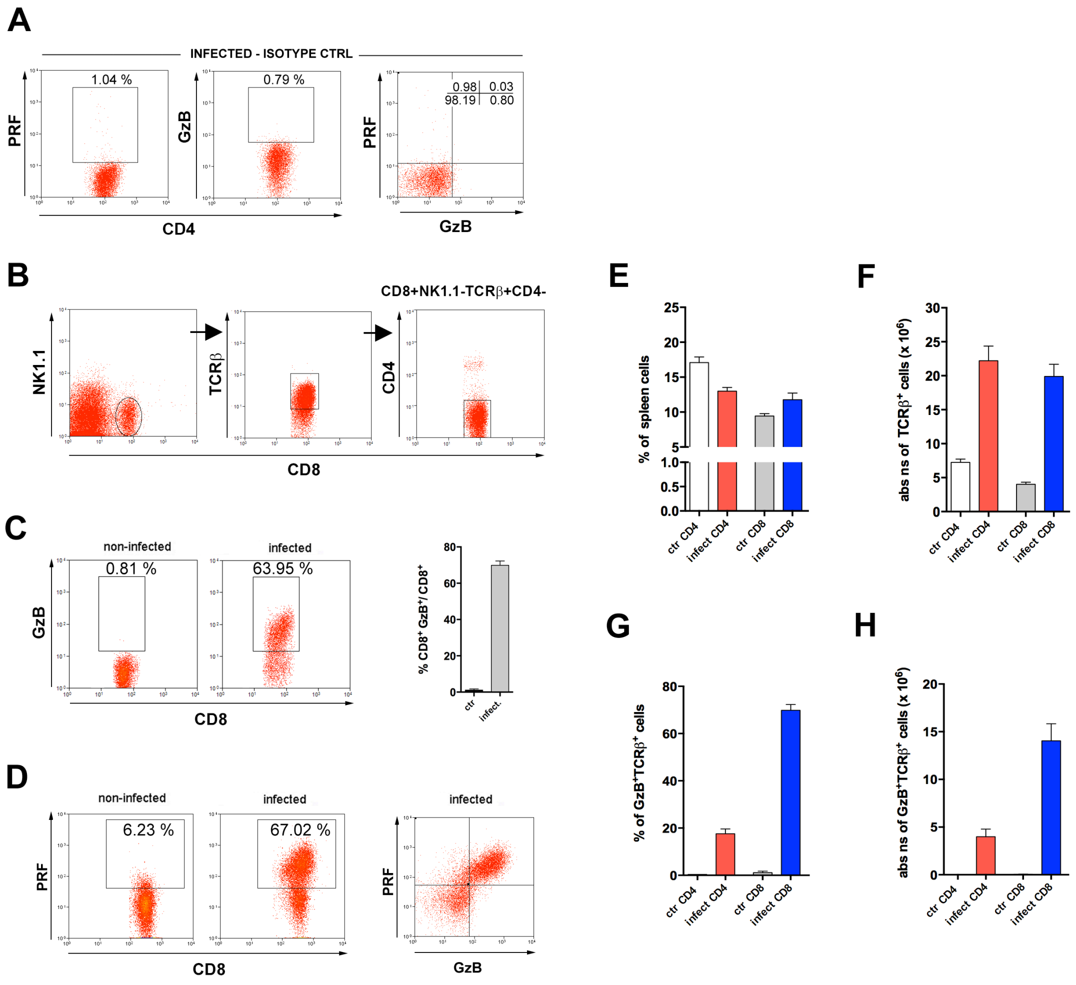


**Figure 1–figure supplement 1: Comparative percentages and absolute numbers of GzB$^+$ and PRF$^+$ T cell subsets. (A)** Isotype control staining of CD4$^+$ T cells, gated as in Fig. 1A, from infected mice. **(B)** Gating strategy for **C** and **D** plots. **(C)** Representative dot plots of GzB staining and mean frequency of GzB$^+$CD8$^+$T cells (on the right). **(D)** Representative dot plots of PRF and GzB staining in CD8$^+$ T cells. **(E)** Mean frequency and **(F)** absolute numbers of CD4$^+$ and CD8$^+$ T cells in the spleen of non-infected (ctr) and infected mice. **(G)** Mean frequency and **(H)** absolute numbers of GzB$^+$CD4$^+$ and GzB$^+$CD8$^+$ T cells. Mice individually analyzed at day 14 pi (n=4); error bars= SEM. Data are representative of 6 independent experiments.

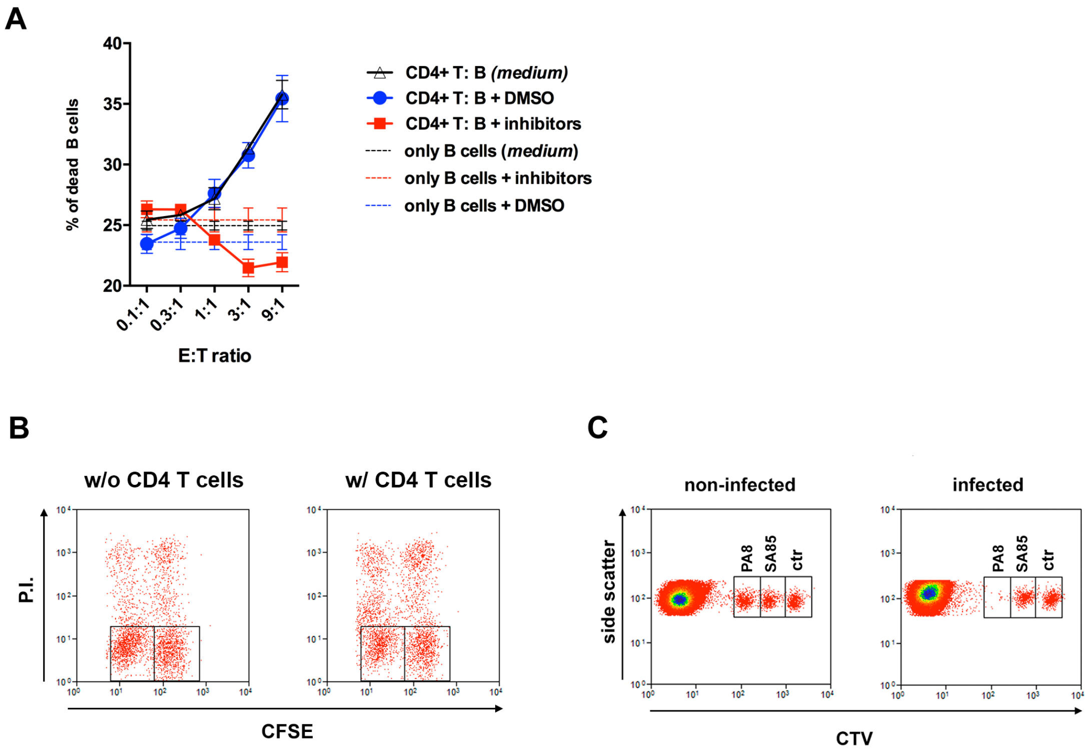


**Figure 2-figure supplement 1: Cytotoxicity assays. (A)** Cytotoxic assays were performed as in **Figure 2A** and in the presence of GzB (Z-AAD-CMK, 10 μM) and PRF (Concamycin A, 0.5 μM) inhibitors (red line and symbols) or DMSO (blue line and symbols). **(B-C)** Gating strategy of cytotoxic assays: **(B)** Gating strategies and PI staining of IC-21 macrophages loaded with total amastigote protein extract ($CFSE^{lo}$) or not ($CFSE^{hi}$) and co-cultured with 4 x $10^5$ purified $CD4^+$ T cells (right), or not (left); specific cytotoxicity values are shown in **Figure 2B**; **(C)** Gating strategies for CTV-stained target splenocytes, loaded with the indicated antigenic peptides or not (ctr), 20 h after i.v. injection in non-infected or Y strain-infected mice. The *T. cruzi*-derived PA8 (class I-restricted) and SA85 (class II-restricted) peptides were employed. Specific cytotoxicity values for the *in vivo* assay are shown in **Figure 2C**.

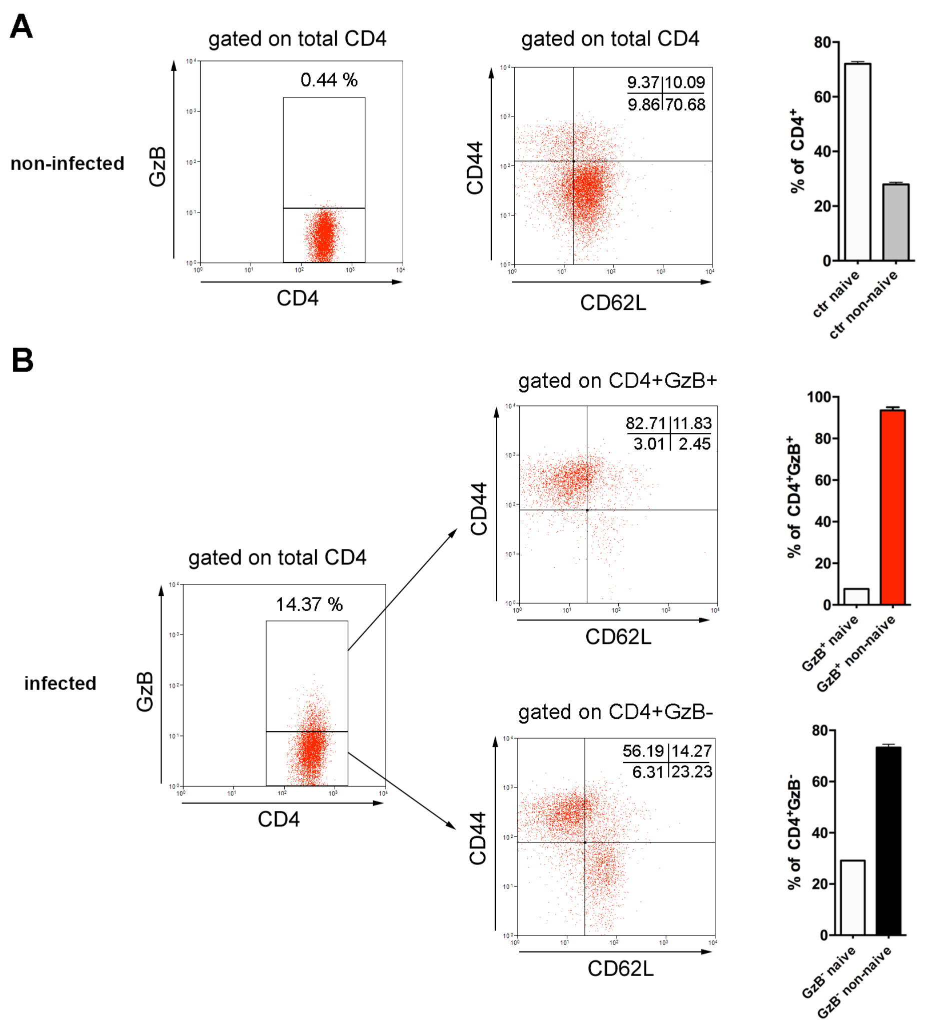


**Figure 2-figure supplement 2: The majority of $GzB^+CD4^+$ and $GzB^-CD4^+$ T lymphocytes are non-naïve (activated effectors and memory) cells.** Representative dot plots and gating strategy of staining for GzB, CD44 and CD62L (on the left) and mean frequency of naive ($CD44^{lo}$) and non-naive ($CD44^{hi}$) $CD4^+$ T cells (on the right) in: **(A)** non-infected mice and gated on total $CD4^+TCR\beta^+$ and **(B)** infected mice and gated on $GzB^+$ (top) or $GzB^-$ (bottom) $CD4^+TCR\beta^+$ cells, at day 14 pi. Bars represent mean values of cell frequency of individually analyzed mice; n=4; Error bars = SEM. Data are representative of 3 independent experiments.

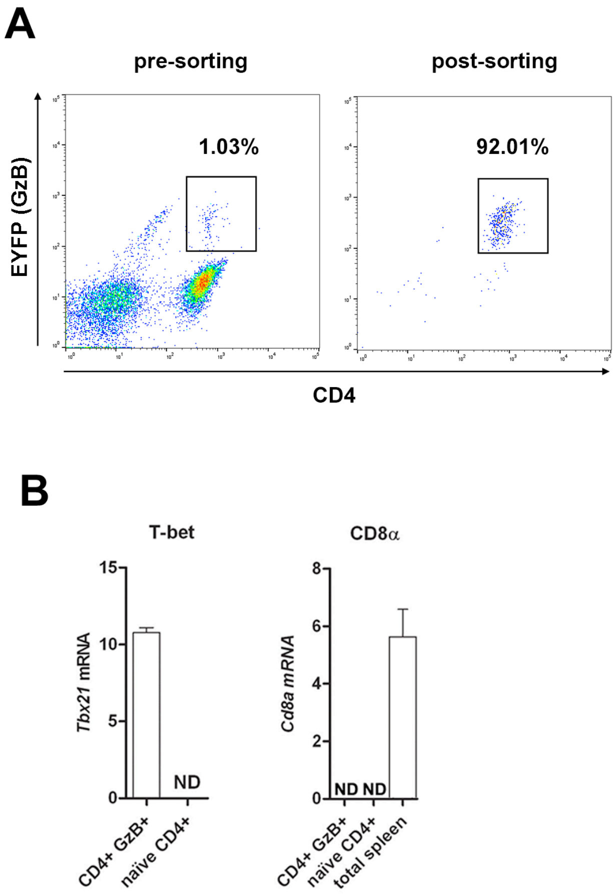


**Figure 2-figure supplement 3: Expression of *Tbx21* and *Cd8a* in sorted GzB$^+$CD4$^+$ T cells. (A)** Sorting gate and frequency of CD4$^+$GzB$^+$ (EYFP)$^+$ T cells obtained from infected and tamoxifen-treated GzmbCreER$^{T2}$/ROSA26EYFP mice. CD4$^+$ cells were enriched by negative selection with magnetic beads, as described in Methods, and then submitted to FACS-sorting. Pre-sorting CD4$^+$ T cell frequency is shown on the left panel and post-sorting on the right. **(B)** Sorted cells had RNA extracted and *Tbx21*, *Runx3d*, *Zbtb7b* and *Cd8a* expression tested by qRT-PCR, as described in Methods. Results for *Runx3d* and *Zbtb7b* expression are shown in **Fig. 2H.** ND = not detected.

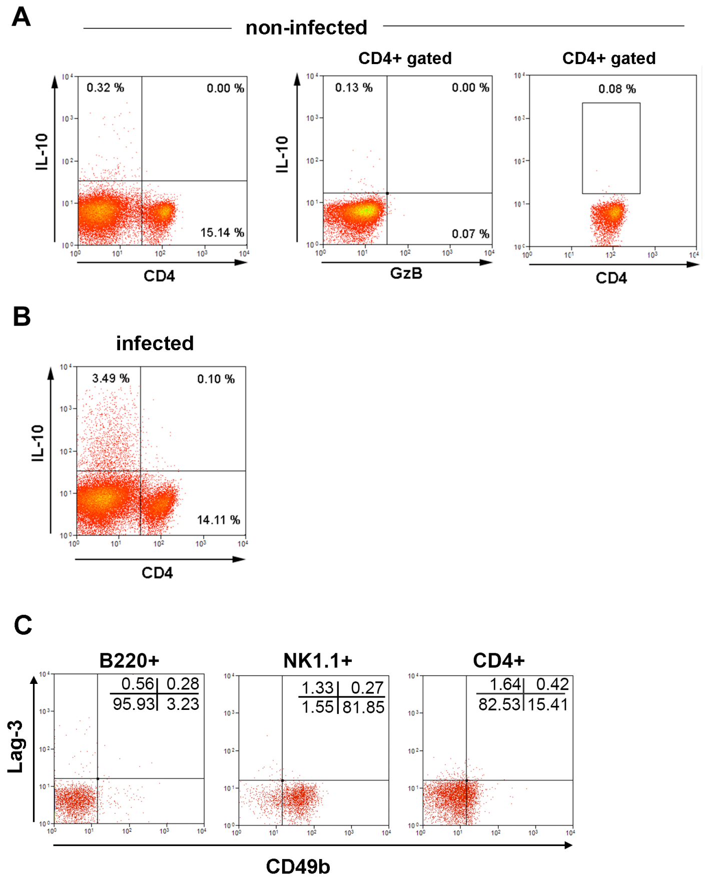


**Figure 3-figure supplement 1: IL-10, Lag-3 and CD49b expression. (A)** Representative dot plots and gating strategy for IL-10 expression. Total spleen cells (left) and gated $CD4^+$ T cells (middle and right panels) of non-infected control mice. **(B)** Representative dot plot of IL-10 staining in total splenocytes of infected mice. Dot plot and mean frequencies of IL-$10^+$ cells among $CD4^+GzB^-$ and $CD4^+GzB^+$ T cells in infected mice are shown in **Fig. 3A. (C)** Representative dot plot of Lag-3 and CD49b expression on gated $B220^+$, $NK1.1^+$ or total $CD4^+$ T cells from the spleen of non-infected mice (left, middle and right panels, respectively). Mean frequencies of Lag-$3^+CD49b^+$ and Lag-$3^+CD49b^-$ cells among $CD4^+GzB^-$ and $CD4^+GzB^+$ T splenocytes of infected mice are shown in **Fig. 3C.** Data are representative of 2 independent experiments.

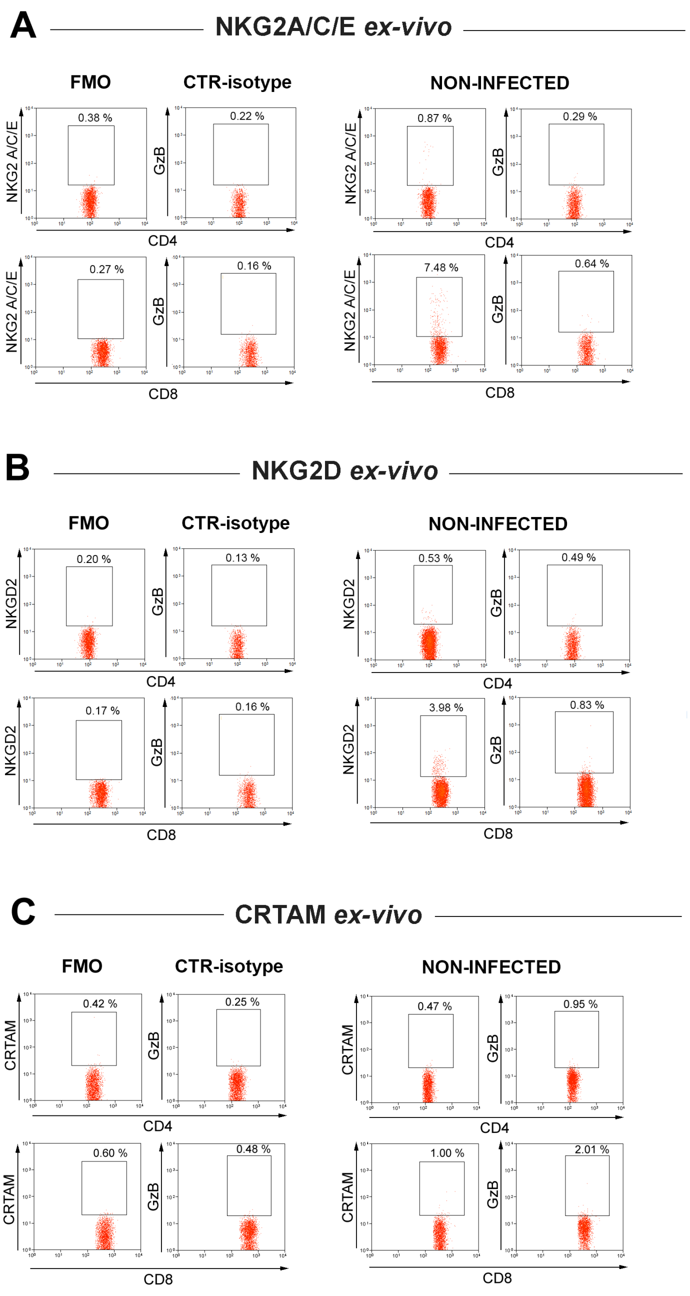


**Figure 3-figure supplement 2: Representative dot plots and gating strategies for NKG2A/C/E, NKG2D and CRTAM staining on CD4$^+$ and CD8$^+$ GzB$^+$ T cells.** Staining of spleen cells from non-infected controls (right panels) and FMO and isotype controls of spleen cells from B6 infected mice (left panels) are shown. Representative dot plots and mean frequencies of CD4$^+$GzB$^+$ and CD8$^+$GzB$^+$ T cells expressing these cytotoxic markers are shown in **Figure 3F-H** and Figure 3-figure supplement 3, respectively.

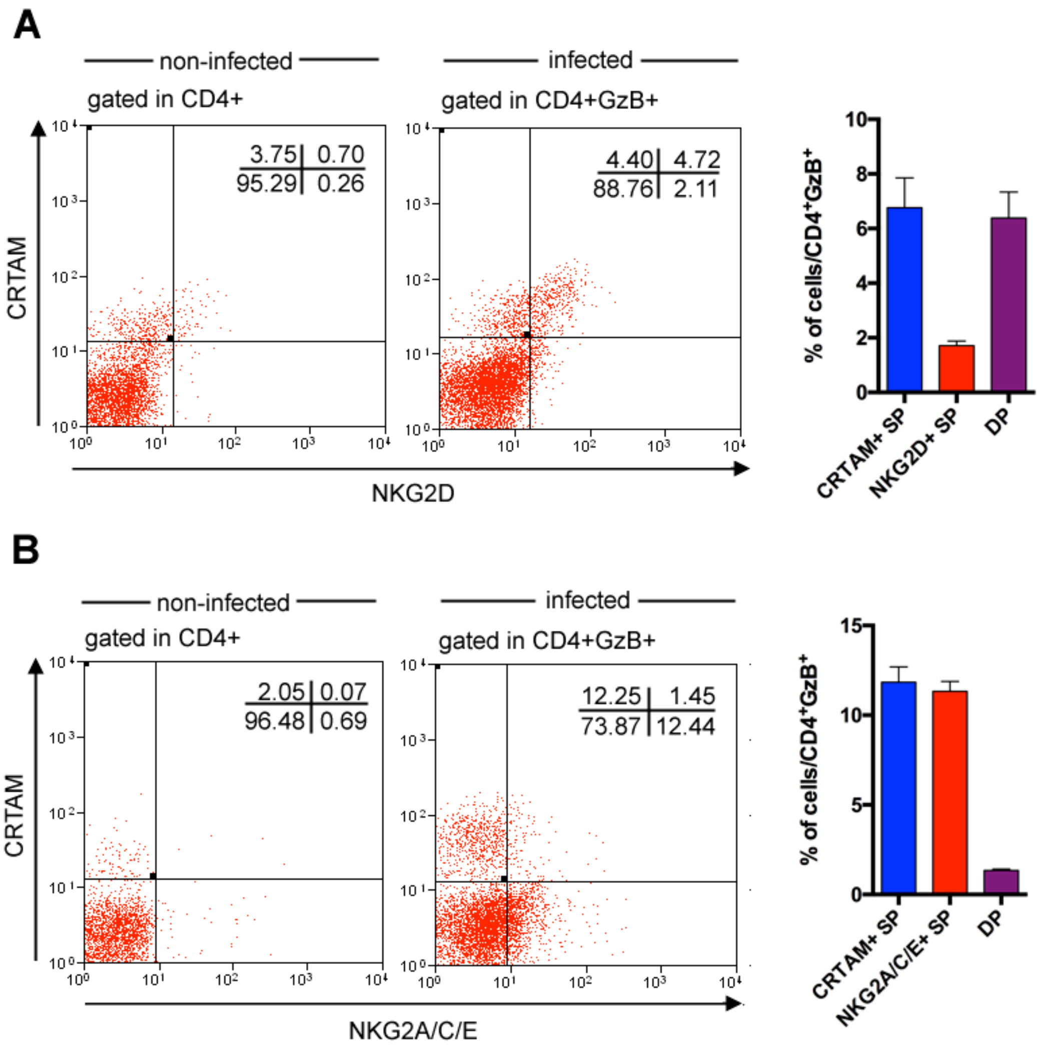


**Figure 3-figure supplement 3: Most $CD4^+GzB^+NKG2D^+$ T cells express the CRTAM cytotoxic marker while only a minority of $CD4^+GzB^+$ T cells co-expresses CRTAM and NKG2A/C/E molecules. (A and B)** Representative dot plots of CRTAM and NDG2D **(A)** or CRTAM and NKG2A/C/E **(B)** staining of spleen cells from non-infected and infected mice, gated on total $CD4^+$ or on $CD4^+GzB^+$ T cells, respectively, as indicated. Bars graphs on the right represent mean values of individually analyzed infected mice; n= 6; error bars = SEM; Data are representative of 3 independent experiments.

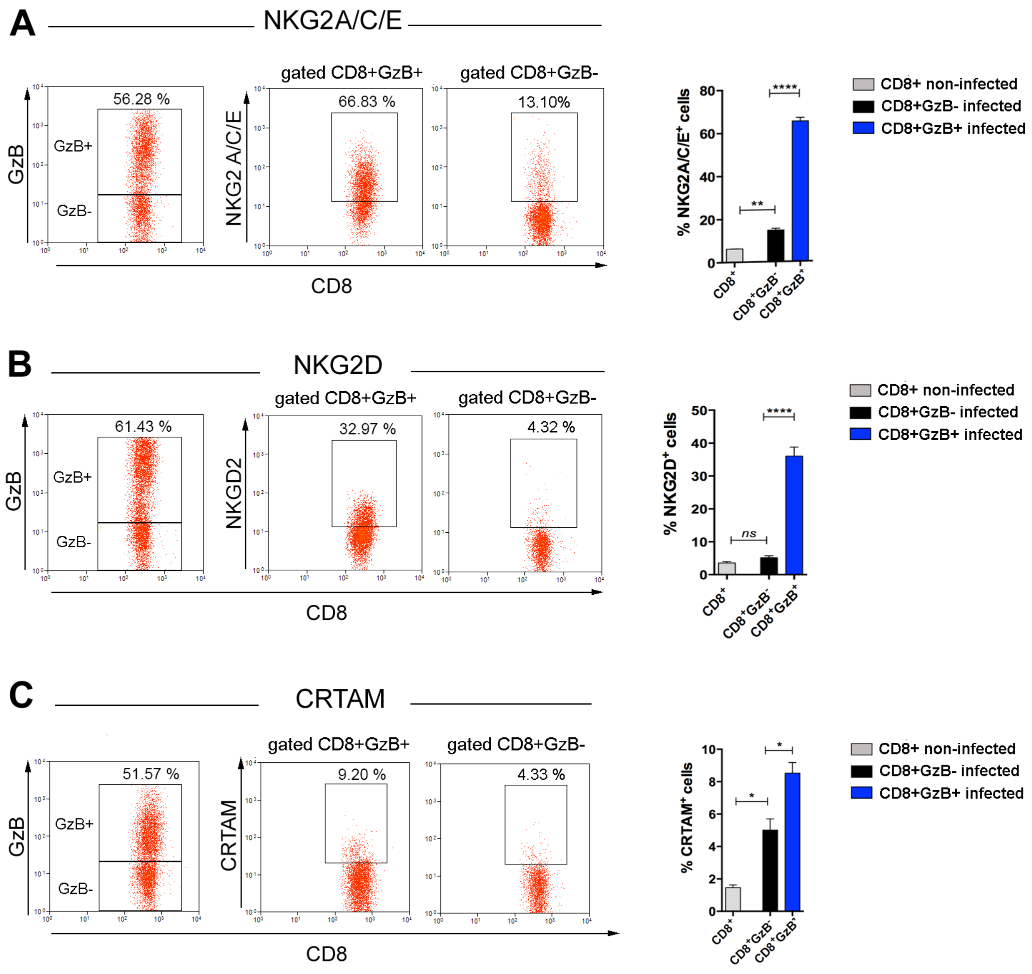


**Figure 3-figure supplement 4: Expression of NKG2A/C/E, NKG2D and CRTAM cytotoxic markers by CD8$^+$GzB$^+$ T cells.** Representative dot plots and gating strategies of spleen cells from B6 infected mice (left panels). Graphs on the right: bars represent mean values of cell frequency of each indicated cell subpopulation in individually analyzed mice; n=4; Error bars = SEM. Data are representative of 3 independent experiments.

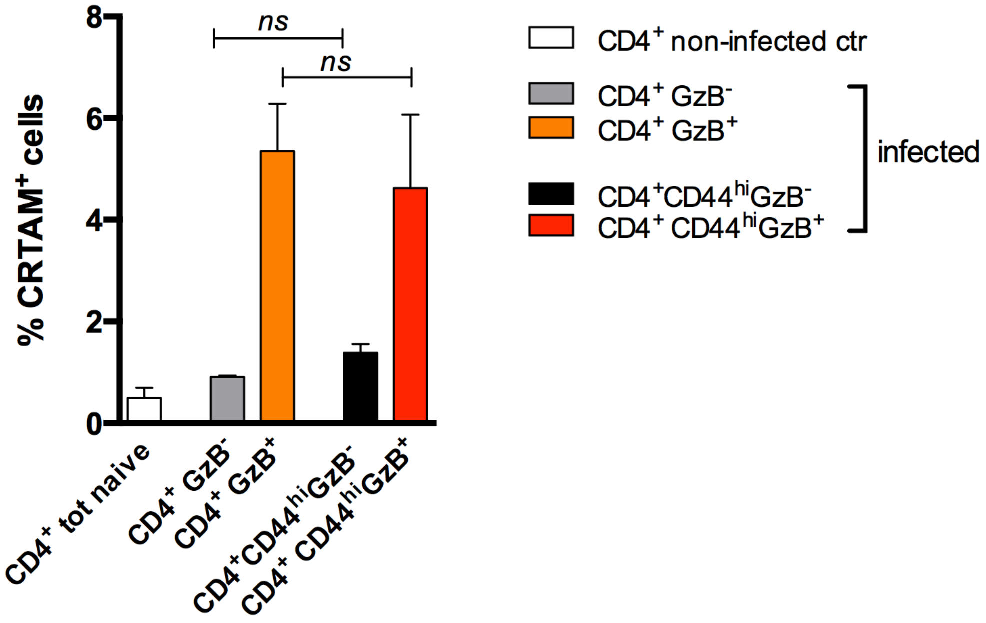


**Figure 3-figure supplement 5: Frequency of CRTAM-expressing cells among splenic GzB$^+$ and GzB$^-$ CD4$^+$ T cells, gating or not on CD44$^{hi}$ cells.** Bars represent mean values of the frequency of CRTAM$^+$ cells among: CD4 naïve cells of non-infected (ctr) mice (white bar) and CD4$^+$GzB$^-$ (gray bar), CD4$^+$GzB$^+$ (orange bar), CD4$^+$CD44$^{hi}$GzB$^-$ (black bar) or CD4$^+$CD44$^{hi}$GzB$^+$ (red bars) of infected mice at day 14 pi; Gating as shown in **Figure 1A-B, Figure 3H and Figure 7-figure supplement 2.** Mice were individually analyzed, n= 4; Error bars= SEM; *ns*= non-significant, Student t test.

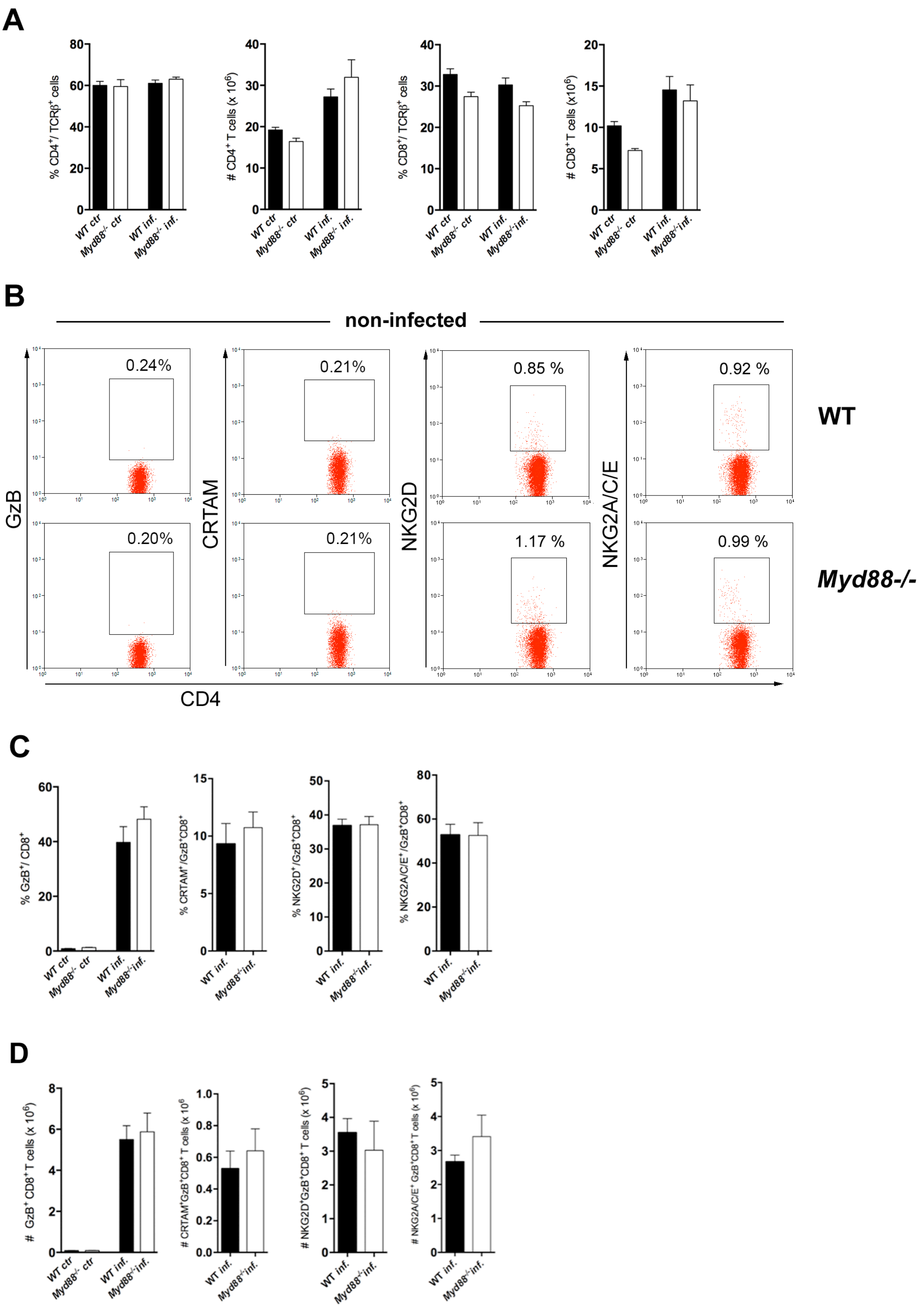


**Figure 4-figure supplement 1: Equivalent frequencies and absolute numbers of total CD4$^+$ T cells in the spleens of infected *Myd88*$^{-/-}$ and WT mice. (A)** Mean frequencies and absolute numbers of total CD4$^+$ and CD8$^+$ T cells in the spleens of non-infected (ctr) and infected *Myd88*$^{-/-}$ and WT (B6) mice, at day 13 pi. **(B)**

Representative dot plots and gating strategies for GzB, CRTAM, NKG2D and NKG2A/C/E staining of spleen cells from non-infected (ctr) WT (B6) and *Myd88*$^{-/-}$ mice. Staining results obtained with infected mice are shown on **Figures 4C-4E**. **(C)** Mean frequencies and **(D)** absolute numbers of GzB$^{+}$, GzB$^{+}$CRTAM$^{+}$, GzB$^{+}$NKG2D$^{+}$ and GzB$^{+}$NKG2A/C/E$^{+}$ cells among CD8$^{+}$ T cells in the spleen of WT and *Myd88*$^{-/-}$ infected mice at day 13 pi, gated as in Sup. Fig. 7. Bars represent mean values in each group; error bars = SEM; n= 5 individually analyzed mice. Results are representative of 2 independent experiments.

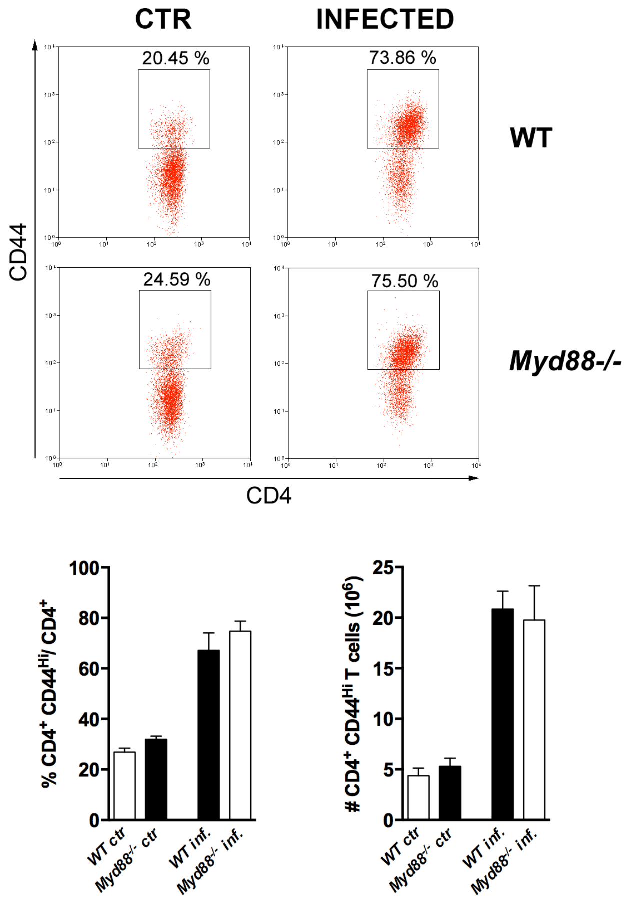


**Figure 4-figure supplement 2: Equivalent frequency and absolute number of activated CD4$^+$ T cells in WT and *Myd88$^{-/-}$* mice infected with *T. cruzi.*** Representative dot plots (top) and mean frequencies and absolute numbers (bottom) of CD4$^+$CD44$^{hi}$ T cells in the spleens of infected WT (B6) and *Myd88$^{-/-}$* mice, at day 14 pi. Mice were individually analyzed; n=4; error bars = SEM. Representative of 2 independent experiments.

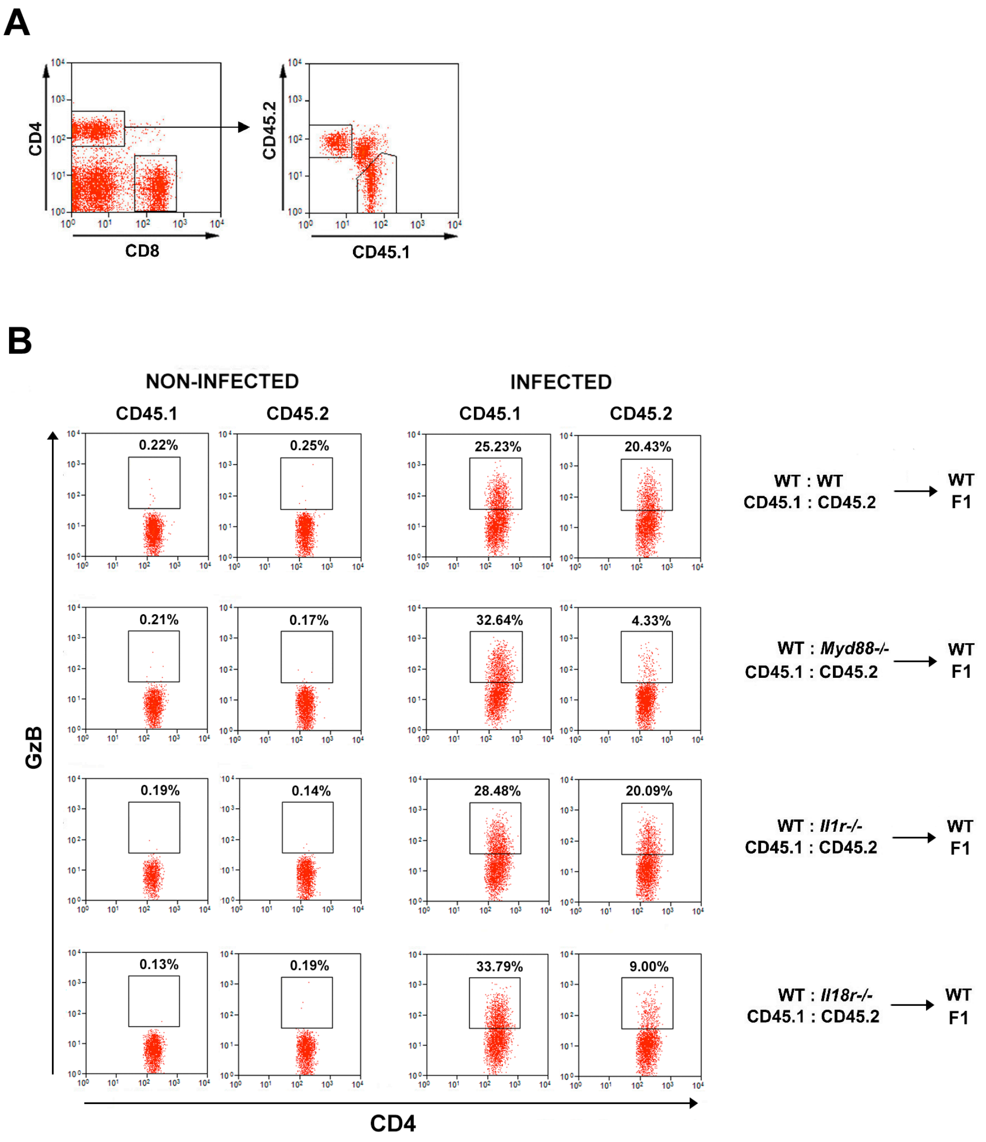


**Figure 5-figure supplement 1: GzB expression in CD4$^{+}$ T cells of mixed-BM chimeric mice.** Gating strategy and representative dot plots of **(A)** CD45.1$^{+}$ and CD45.2$^{+}$ cells (on the right) gated CD4$^{+}$CD8$^{-}$ T lymphocytes (on the left) from the spleens of mixed-BM chimeric mice. **(B)** GzB expression in CD4$^{+}$CD45.1$^{+}$ (WT) and CD4$^{+}$CD45.2$^{+}$ (WT or KO) T cells from the spleens of non-infected and infected chimeric mice at day 14 pi, gated as in **(A).** Mean frequencies and absolute numbers of CD4$^{+}$GzB$^{+}$ in each group are shown in **Figure 5F**.

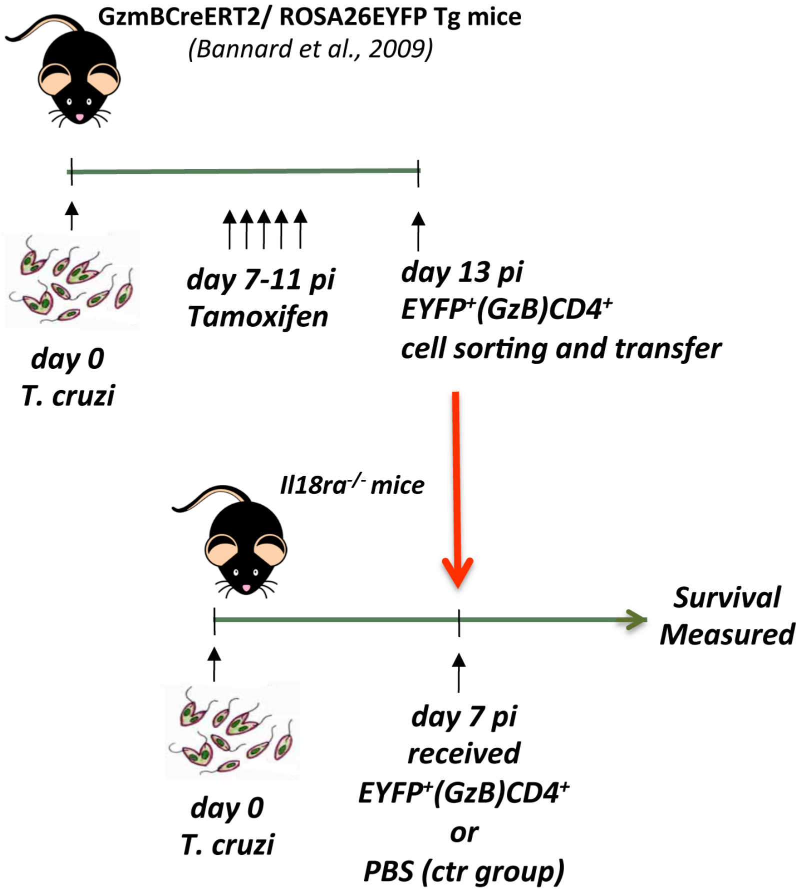


**Figure 6-figure supplement 1: Time-line of the adoptive transfer experiment shown on Figure 6G.** Description of the protocol is available in the M&M section.

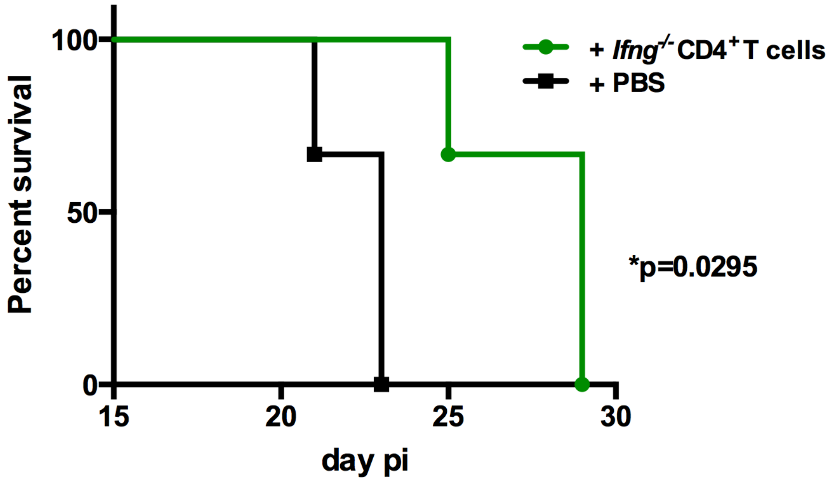


**Figure 6-figure supplement 2: The adoptive transfer of *Ifng*$^{-/-}$ CD4$^{+}$ T cells increased survival to infection.** Survival curve of *Il18ra*$^{-/-}$ mice, infected with 1 x $10^3$ trypomastigotes of the Y strain and adoptively transferred with sorted (>98% pure) *Ifng*$^{-/-}$ CD4$^{+}$ T cells (green curve) or not (PBS group, black curve); n=6 mice in each group; p= 0.0295 Log-rank (Mantel-Cox) test.

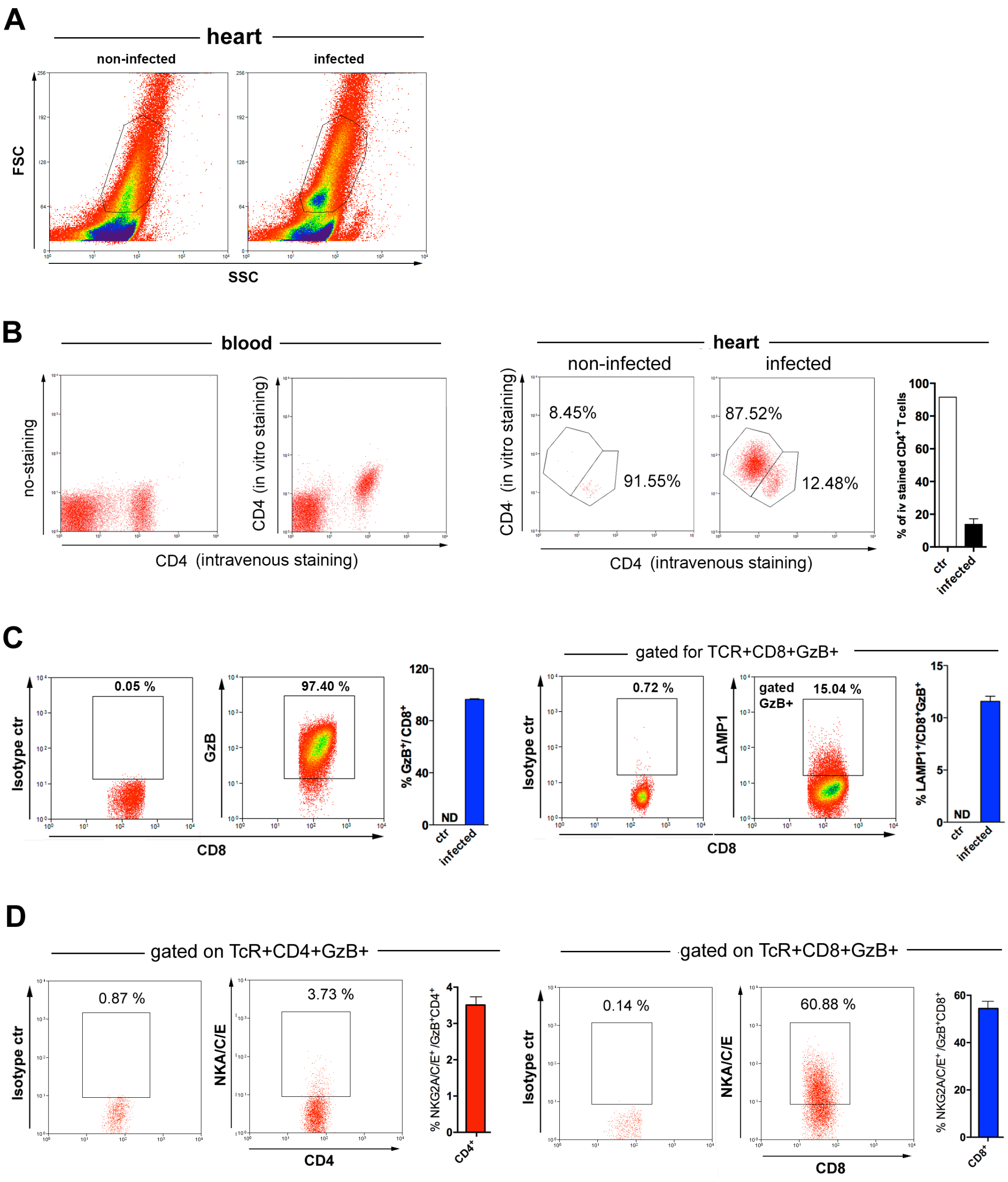


**Figure 7-figure supplement 1: CD4$^+$ and CD8$^+$ T cells in the heart of mice infected with *T. cruzi*.** **(A)** Representative FSC x SSC dot plots and gating strategy of cells from the hearts of non-infected (left panel) and infected (right panel) WT mice, employed for data analysis shown on this figure and Figure 7. **(B)** Representative dot plots of blood samples from infected mice injected iv with anti-CD4-FITC mAb, 3 min before euthanasia, with no other staining (left panel) or after staining with anti-

CD4-PECy7 mAb *in vitro* (right panel). Representative dot plots of cells from the heart of non-infected and infected mice, injected *iv* with anti-CD4-FITC mAb and stained *in vitro* with anti-CD4-PECy7 (two panels on the right), and mean frequencies of intravenously stained $CD4^+$ T cells in the hearts of non-infected and infected mice. Mice were individually analyzed; n=3; error bars = SEM. Representative of 2 independent experiments. **(C)** Representative dot plots and gating strategy of isotype and GzB (left panels) or LAMP1 (right panels) staining and corresponding mean frequencies of $GzB^+CD8^+$ and $LAMP1^+CD8^+$ cells among gated $CD8^+$ T cells infiltrating the infected myocardium at day 14 pi. **(D)** Representative dot plots of isotype and NKG2A/C/E staining on gated $CD4^+GzB^+$ (left panels) or $GzB^+CD8^+$ (right panels) T cells and corresponding mean frequencies of $NKG2A/C/E^+$ cells among $GzB^+CD4^+$ and $GzB^+CD8^+$ T cells infiltrating the infected myocardium at day 14 pi. Individually analyzed mice, n= 4; error bars= SEM; ND= not detected. Data are representative of 2 independent experiments.

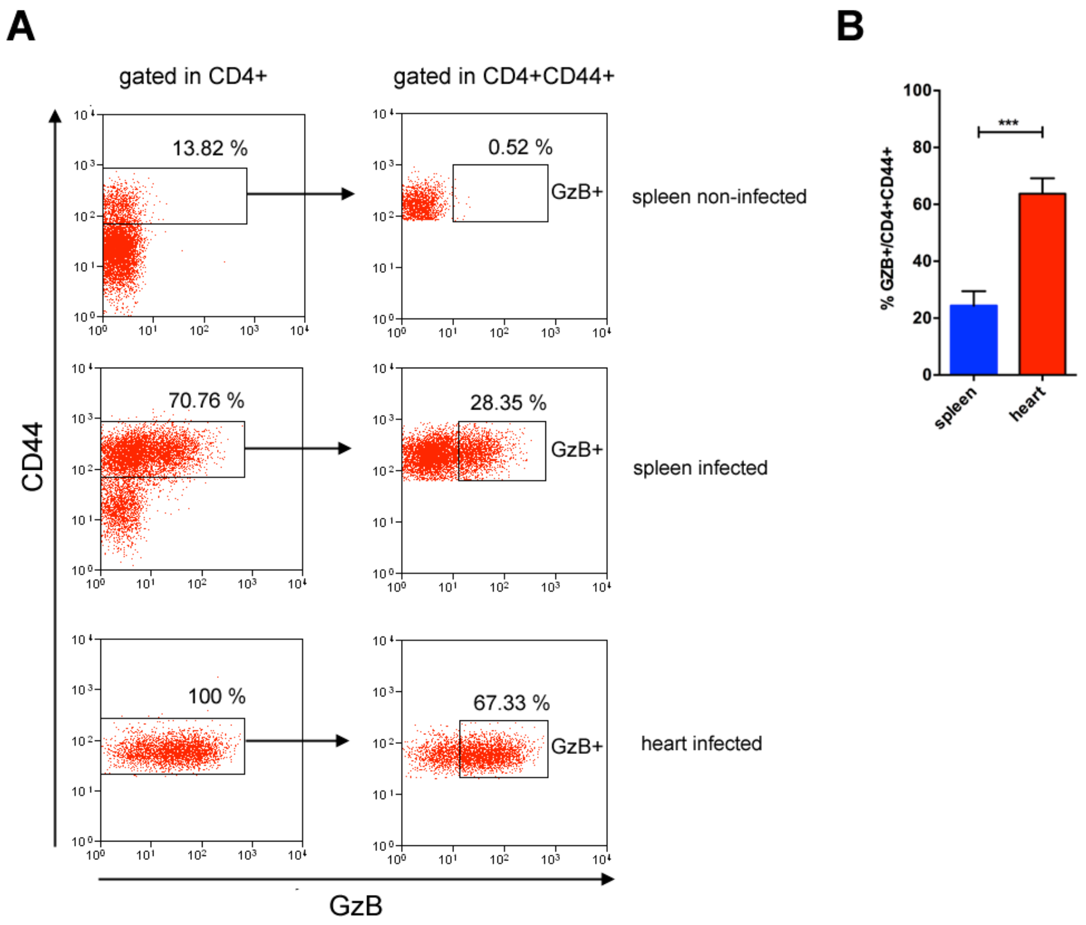


**Figure 7-figure supplement 2: GzB$^+$CD4$^+$ T cells are enriched among activated/memory CD4$^+$ T cells in the heart compared to the spleen of mice infected with *T. cruzi.*** **(A)** Representative dot plots of GzB and CD44 staining of CD4$^+$ T cells in the spleen and in the heart, gated as in Figures 1 and 7A. **(B)** Frequencies of GzB$^+$ cells among activated/memory CD4$^+$ T cells in the spleen and in the heart of individually analyzed infected mice; n= 9; error bars= SEM, $^{***}p \leq 0.001$ Student t test. Data are compiled from three independent experiments.

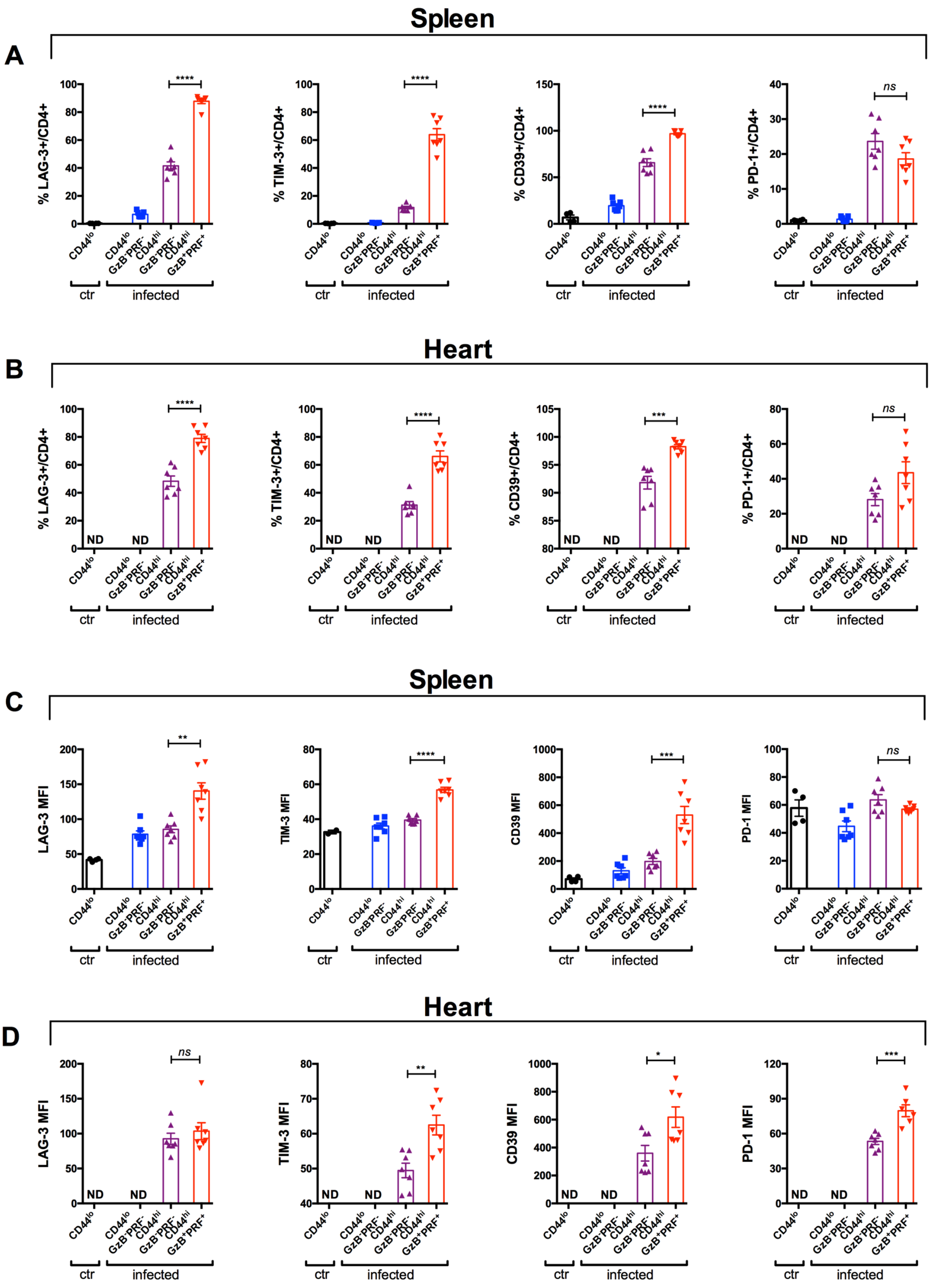


**Figure 7-figure supplement 3: Cytotoxic CD4+ T cells express the higher levels of LAG-3, TIM-3, CD39 or PD-1 molecules. (A and B)** Frequency of cells expressing LAG-3, TIM-3, CD39 or PD-1 molecules among naïve cells form non-infected mice ($CD44^{lo}$, ctr) and naïve cells ($CD44^{lo}GzB^{-}PRF^{-}$), activated non-cytotoxic

($CD44^{hi}GzB^{-}PRF^{-}$) or activated cytotoxic ($CD44^{hi}GzB^{+}PRF^{+}$) cells from infected mice, and **(C and D)** their respective levels of expression (mean fluorescence intensity, MFI), in the spleen and in the heart at day 14 pi, as indicated. Bars represent mean frequency **(A and B)** or MFI (**C and D)** ± SEM of individually analyzed mice; n=7; *ns*= non-significant, $^{*}p \leq 0.05$, $^{**}p \leq 0.01$, $^{***}p \leq 0.001$, $^{****}p \leq 0.0001$ Student t test; ND= not detected. Data are compiled from 2 independent experiments.

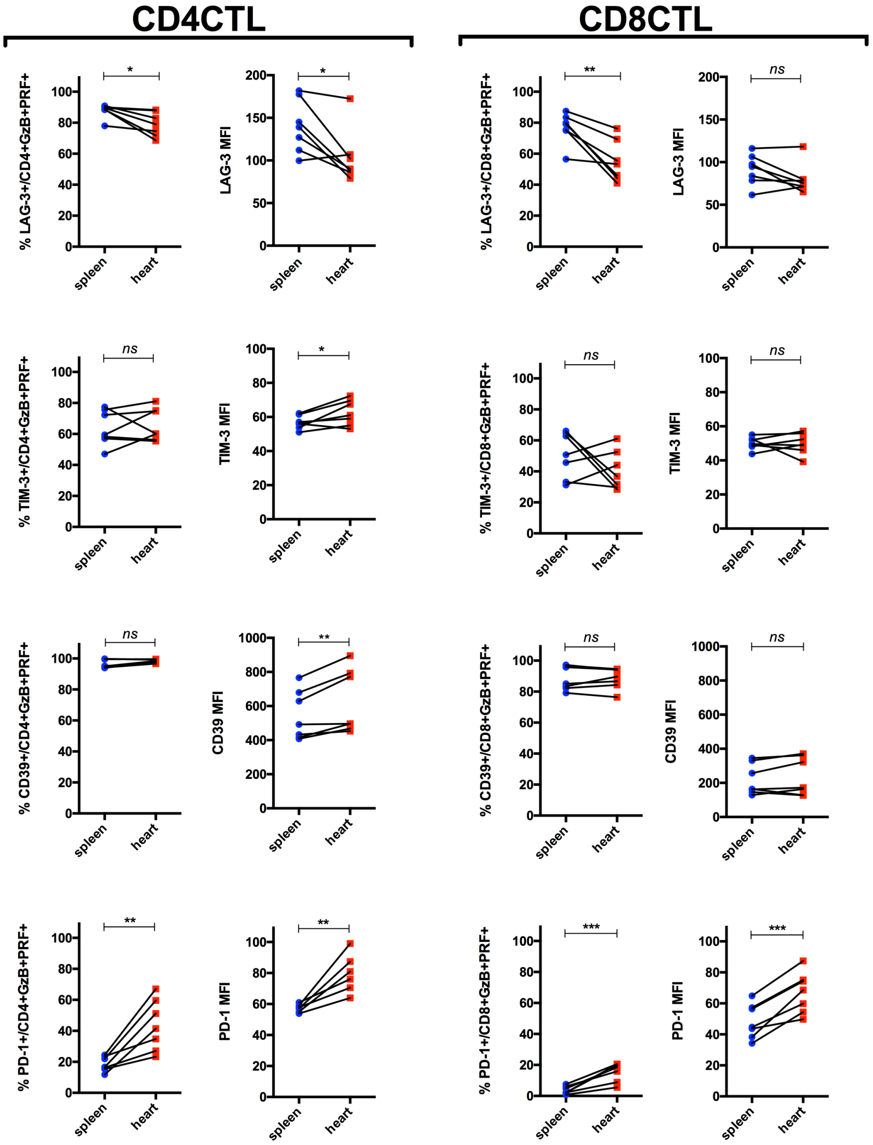


**Figure 7-figure supplement 4: The expression of PD-1, CD39 and Tim-3, but not of Lag-3, is increased on CD4CTLs infiltrating the heart, compared to their levels in the spleen.** Graphs indicate the frequency of $GZB^{+}PRF^{+}$ T cells expressing Lag-3, Tim-3, CD39 and PD-1 among $CD4^{+}$ and $CD8^{+}$ T cells (CD4TLs on the left and CD8CTLs on the right), and their corresponding MFI, in the spleen (blue dots) and in the heart (red squares), at day 14 pi. Symbols represent individually analyzed mice; n = 7; *ns*= non-significant, $^{*}p \leq 0.05$, $^{**}p \leq 0.01$, $^{***}p \leq 0.001$; paired Student t test. Data are compiled from 2 independent experiments.

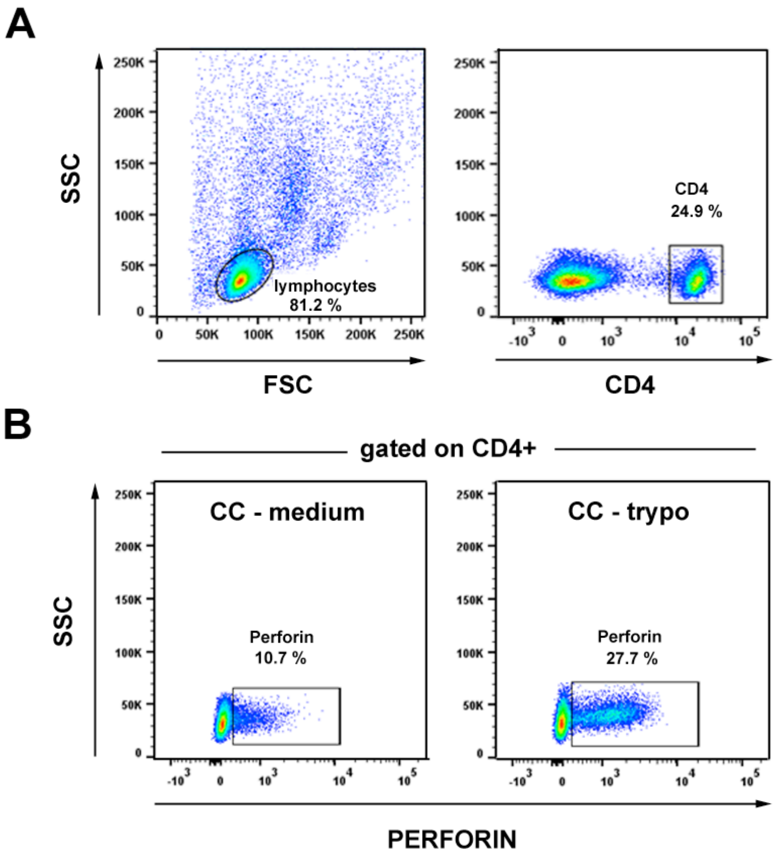


**Figure 7-figure supplement 5: CD4$^+$ T cells expressing PRF are found in the circulation of patients suffering from chronic Chagas cardiomyopathy (CC). (A)** Dot plot showing the gating strategy for CD4$^+$ T cell staining on PBMC. **(B)** Gating strategy and representative dot plots of PRF staining, gated CD4$^+$ T cells as in (A), from CC patient incubated with medium (on the left), or with trypomastigote antigens (trypo) (on the right). Frequencies of PRF$^+$CD4$^+$ T cells and MFI of PRF staining for CC patients and HD controls are shown on **Figures 7G-7I**.

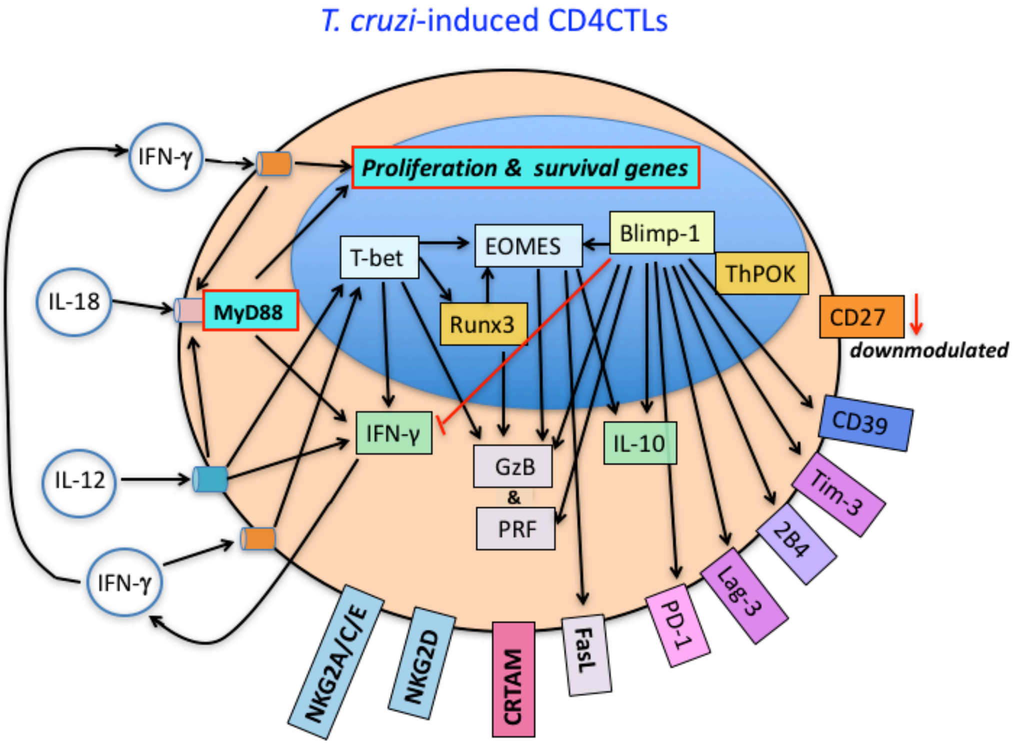


**Figure 7-figure supplement 6: Predicted model for the generation of CD4CTLs cells in response to *T. cruzi* infection.** This cartoon summarizes our results and hypothesis and also contains information from previous published studies, cited in this work. IL-12 produced during infection acts on TCR-triggered $CD4^+$ T cells, leading to T-bet expression, which promotes IFN-γ upregulation and induces *Runx3d* and *Eomes* gene expression. In turn, IFN-γ- and IL-12-signaling induce IL-18R expression. T-bet, Eomes and Runx3d drive the expression of cytotoxic effector molecules, including GzB, PRF and FasL. IL-10 expression can be induced by Blimp-1 and Eomes, although few $IL-10^+$ $CD4^+GzB^+$ T cells were found in our infection model. As opposed to intraepithelial CD4CTLs of the gut, ThPOK expression is not downmodulated in splenic $CD4^+GzB^+$ T cells, until day 14 pi. Only around 18% of $CD4^+GzB^+$ T cells produce IFN-γ at the peak of the $CD4^+$ T cell response (14 dpi), and IFN-γ is necessary for $CD4^+GzB^+$ T cell proliferation and/or survival. Most $CD4^+GzB^+$ T cells express Lag-3, Tim-3, PD1 and CD39 immunoregulatory molecules, possible under the control of Blimp-1, while CD27 is downmodulated. Notably, the expansion and/or survival of $CD4^+GzB^+$ T cells expressing CRTAM, NKG2D, and NKG2A/C/E cytotoxic markers depend on T-cell intrinsic IL-18R/MyD88 signaling.
